# Early root-root interactions weaken foliar defense responses against Septoria tritici blotch in a durum wheat varietal mixture

**DOI:** 10.1101/2025.07.30.667709

**Authors:** Laura Mathieu, Amandine Chloup, Soline Marty, Justin Savajols, Claudia Rouveyrol, Josep Valls, Pierre Pétriacq, Jean-Benoît Morel, Louis Valentin Méteignier, Elsa Ballini

**Affiliations:** PHIM Plant Health Institute, Univ Montpellier, INRAE, CIRAD, Institut Agro, IRD, Montpellier, France; Univ Bordeaux, INRAE, UMR1332 Biologie du Fruit et Pathologie, 33882 Villenave d’Ornon, France; Bordeaux Metabolome, MetaboHUB, PHENOME-EMPHASIS, Villenave d’Ornon, France; Univ Bordeaux, INRAE, UMR 1366 OENO - Axe MIB, ISVV, 33140, Villenave d’Ornon, France; PHIM Plant Health Institute, Univ Montpellier, CIRAD, Institut Agro, INRAE, IRD, Montpellier, France

**Keywords:** Durum wheat, *Triticum turgidum*, Septoria tritici blotch, varietal mixtures, plant-plant interactions, belowground processes, competition

## Abstract

The interactions between co-cultivated plant cultivars are increasingly recognized as influencing their susceptibility to pathogens in mixtures. However, the underlying mechanisms remain largely unexplored. Using a model of durum wheat cultivar mixtures where susceptibility to Septoria foliar disease is increased, we combined aerial and root phenotyping with transcriptional analyses and untargeted metabolomics to elucidate the potential signaling cascade driving this modulation of susceptibility. We observed contrasting root architectures between cultivars in mixture. Molecular analysis showed a delayed induction of defense-related genes and metabolites following pathogen inoculation in plants grown in mixture compared to pure stand. The findings suggest that root architecture potentially triggers a competitive response that could delay the induction of defense responses following pathogen inoculation. Altogether, these results point to a possible interplay between root architecture, resource competition, plant metabolism, and defense modulation in shaping plant–pathogen interactions within varietal mixtures.

**Summary statement:** In a durum wheat mixture, increased susceptibility to Septoria tritici blotch is linked to delayed defense response induction, likely triggered by early root competition resulting from contrasting root architectures. This reveals a novel root-mediated mechanism by which plant-plant interactions modulate susceptibility to aerial pathogens.

## Introduction

Plant–plant interactions are increasingly recognized as a key factor that can influence plant health, particularly by modulating susceptibility to pathogens, a phenomenon known as Neighbor-Modulated Susceptibility (NMS) (Pélissier *et al*. 2021a). Recent studies have demonstrated that plants can exchange signals that influence immune responses, often resulting in enhanced resistance to pests and diseases (Pélissier, Violle & Morel 2021b; Pélissier *et al*. 2023). Notably, the level of disease susceptibility modulated by plant–plant interactions can be, in some cases, comparable to that conferred by intrinsic basal immunity (Mathieu, Ducasse, Ballini & Morel 2024b). In durum wheat, the mixture of two genotypes, Cultur and Atoudur, was previously described to modulate susceptibility to *Zymoseptoria tritici*, a fungal pathogen responsible for Septoria tritici blotch (STB) (Pélissier *et al*. 2021a). Specifically, in the presence of Atoudur, Cultur becomes more susceptible to STB, but the underlying mechanisms remain unknown.

Plant-plant interactions can be envisioned as a two-step process: first, plants exchange signals, through the soil or air; second, these signals trigger morphological and/or physiological responses in the plant under consideration (later called focal plant). Regarding signaling, direct plant-plant interactions can be mediated by molecular cues or access to resources (Mathieu, Ballini & Morel 2025). A previous study on the Cultur-Atoudur mixture demonstrated that the introduction of a physical barrier between root genotypes disrupted the modulation of susceptibility to leaf rust (Pélissier *et al*. 2021a). This indicated the involvement of belowground interaction, though the underlying mechanism remains unknown. Belowground direct plant-plant interactions are often related to root exudates (Mathieu, Ballini, Morel & Méteignier 2024a) and/or nutrient availability (Craine & Dybzinski 2013). However, the belowground signals by which plants influence one another, especially within varietal mixtures, are still poorly understood. In the case of the Cultur–Atoudur mixture, further investigation is needed to elucidate whether changes in disease susceptibility are driven by root exudates, which could be examined by placing exudate-capturing barriers between roots, or by differential access to nutrients, assessable for instance through root architecture measurements.

The modulation of susceptibility by plant–plant interactions may arise from two key response mechanisms. First, plant-plant interactions can influence leaf morphological traits, which may, in turn, affect plant susceptibility. Indeed, traits such as plant height and heading date are negatively correlated with resistance to STB; i.e., STB-resistant cultivars tend to be shorter and exhibit earlier heading (Camacho-Casas, Kronstad & Scharen 1995). Notably, overlapping genetic loci that influence both heading date and disease resistance have been identified (Gerard, Börner, Lohwasser & Simón 2017). Second, plant–plant interactions may alter immune responses, leading to changed susceptibility as observed in the case of leaf rust pathogen in wheat (Pélissier *et al*. 2021a). More generally, wheat-STB interactions are associated with many molecular changes that have been largely described (Brennan, Benbow, Mullins & Doohan 2019). Investigating whether plant–plant interactions influence these molecular changes could enhance the yet limited molecular understanding of susceptibility modulation in mixtures (Subrahmaniam *et al*. 2018). To test these hypotheses on the Cultur-Atoudur mixture, aerial traits could be measured to assess morphological changes, and gene expression and metabolite accumulation could be analyzed to investigate potential immune modulation.

In this study, we investigated how plant-plant interactions within wheat varietal mixtures influence susceptibility to STB. The objectives were to (i) further identify the possible belowground processes that modulate susceptibility, (ii) characterize the morphological and molecular changes associated with responses to neighboring plants, and (iii) determine how plant-plant interactions alter molecular responses to STB infection. Using the model of durum wheat cultivar mixtures between Cultur and Atoudur, we combined aerial and root phenotyping with transcriptional analyses and untargeted metabolomics to propose a potential mechanistic framework underlying the modulation of susceptibility.

## Materials and methods

### Plant material

The study utilized two varieties of durum wheat susceptible to Septoria tritici blotch (STB): Cultur and Atoudur, obtained from RAGT in 2007 and 2010, respectively. The lines were selected for their difference in STB susceptibility between pure and mixed conditions (Pélissier *et al*. 2021a).

### Plant growth conditions

The wheat plants were cultivated in a greenhouse at PHIM (Montpellier, France) under controlled conditions. The photoperiod was 16 hours of light and 8 hours of darkness, with a temperature of 24°C/20°C and a PAR intensity of 250 μmol/s/m2.

Plants were grown in 8cm × 8cm × 8cm pots filled with a soil mixture consisting of 50% topsoil, 50% Neuhaus N2 soil, and 133.3 g of TopPhos for 100L soil. Each pot contained two rows: one with four plants of the focal genotype, on which phenotype is observed, and the other with four plants of the neighboring genotype.

For root phenotyping, plants were cultivated in rhizoboxes (20 cm × 40 cm × 2 cm) filled with the same soil mixture, enabling detailed observation of root architecture. The rhizoboxes were inclined at a 30° angle relative to the horizontal plane to encourage root development along the lower face of the box. The sides of the rhizoboxes were shielded from light to prevent unwanted light exposure. In each rhizobox, two plants of the same genotype (pure) or two plants of different genotypes (mixture) were sown against the glass panel, with a spacing of 5 cm between them.

### Inoculation and symptom assessment

The Cultur genotype, either grown in pure stands or mixed with the Atoudur genotype, was cultivated under control conditions or with either an empty porous barrier or the same porous barrier containing polyvinylpolypyrrolidone (PVPP) to block the exchange of chemical molecules, particularly phenolics, between genotypes (Durán-Lara *et al*. 2015). The Cultur genotype was inoculated with the *Zymoseptoria tritici* strain “P1A” (Ballini *et al*. 2020) by applying an inoculum of 10□spores/mL to the last ligulated leaf 21 days after germination using a brush. Following inoculation, plants were placed in transparent plastic bags for three days to maintain high humidity and promote pathogen growth. 17 days post-inoculation, STB symptoms were assessed using an Epson Perfection V370 Photo scanner. The SeptoSympto tool (Mathieu *et al*. 2024c) was used to quantify the surface area of necrosis, which refers to the lesions.

### Root and aerial trait assessment

Root and aerial traits were evaluated at different growth stages under both pure and mixed conditions (Supplementary Fig. S1). Root traits for the Cultur and Atoudur genotypes were assessed 21 days after germination, at the 3-leaf stage allowing the root disentanglement. Roots were thoroughly washed before being scanned using WinRhizo software. Root parameters including root volume, diameter, and the number of forks were quantified. Aerial traits were measured 38 days after germination, focusing on the Cultur genotype in both pure and mixed conditions. Plant height, leaf number, and tiller number were scored. Chlorophyll content was also assessed using a Dualex sensor, providing insights into photosynthetic potential and plant health. Additionally, the flowering rate at 56 days after germination was scored on the Cultur genotype in pure and mixture.

### Sample collection

Leaf samples were collected from the genotype Cultur, with each sample comprising a pool of the last expanded leaves from three different pots (12 leaves total per sample). Root samples were taken from the same three pots. After grinding, samples were divided for transcriptional and metabolomic analyses.

Leaf samples were collected at 10 and 21 days after germination (day of inoculation) from Cultur grown in pure stands or mixed with Atoudur, and at 7 and 14 days post-inoculation from Cultur grown in pure stands or mixed with Atoudur under mock or inoculated treatments (Supplementary Fig. S1). Root samples were collected at 21 days after germination (day of inoculation) from Cultur grown in pure stands or mixed with Atoudur (Supplementary Fig. S1). For leaves, the central sections were cut into 0.5 cm pieces, while for roots, the tips were cut into 0.5 cm sections. The samples were immediately stored in liquid nitrogen.

### Primer design for the high-throughput RT-qPCR chip

A systematic literature review was conducted to identify gene biomarkers of 22 physiological processes and 41 sub-processes (Supplementary Fig. S2, Supplementary Table S1). If available, we preferably selected studies showing gene transcriptional regulation (by RT-qPCR, microarray or RNA-seq) in the wheat leaf of young seedlings. When we could not find studies reporting transcriptional regulation of genes under one given process, we selected studies showing the implication of genes using mutant approaches. If available, we selected already designed PCR primers based on our bibliography analysis. However, if no primers were already published for the gene identified, we designed our own primers. The design of primers followed a set of stringent criteria, as generally suggested in RT-qPCR protocols (e.g. PrimerExpress Software v2.0 Application Manual, Applied Biosystems). To minimize the risk of amplifying contaminating genomic DNA, primers spanning at least one exon-exon junction, or annealing to different exons, were designed when possible. The specificity of each primer was confirmed by comparing its sequence with all predicted wheat coding sequences (CDS) using the Primer3Plus software to ensure that at least one primer of each pair targets a unique site within the set of predicted wheat CDS. Primers specificity and amplicon length were checked for each pair of primers using MFEprimer3.1 and verification of hairpin and homo/heterodimerization was checked with idtdna OligoAnalyzer.

### RNA extraction

Total RNA was extracted using the protocol described in Delteil et al. 2012. Initially, frozen leaf tissues were ground in liquid nitrogen. Cells were lysed by adding 1 mL of Tri-reagent to the powdered sample. Nucleic acids were then separated by adding 200 µL of chloroform, incubating for 10 minutes at room temperature, followed by centrifugation at 13000g for 15 minutes at 4°C. The upper aqueous phase was collected, and 560 µL of isopropanol was added to precipitate the nucleic acids overnight at -80°C, followed by centrifugation at 13000g for 20 minutes. The resulting pellet was washed twice with 500 µL of 70% ethanol and centrifuged at 13000g for 5 minutes at 4°C, and then air-dried for 10 minutes before resuspension in 50 µL of ultrapure water on ice for 1 hour. DNase treatment was performed on 50 µL of RNA using the TermoFisher kit, followed by purification. 200 µL of RNA was mixed with 200 µL of phenol:chloroform:alcohol (25:24:1), and centrifuged at 16,000g for 10 minutes at 4°C. The upper phase was collected and precipitated with 18 µL of 3M sodium acetate and 495 µL of pure ethanol at -80°C for 1 hour, followed by centrifugation at 16000g for 30 minutes. The pellet was washed with 1 mL of 70% ethanol, centrifuged for 5 minutes, dried, and resuspended in 50 µL of ultrapure water. RNA quality and quantity were assessed using a Nanodrop spectrophotometer and verified for integrity via 2% agarose gel electrophoresis. The absence of DNA contamination was confirmed by qPCR using the Promega kit with EGM34 and EGM35 primers. For cDNA synthesis, RNA was diluted to 625 ng/µL in 8 µL, combined with 2 µL oligo(dT) and 3 µL H_2_O, and incubated at 95°C for 5 minutes. Subsequently, 4 µL 5X buffer, 2 µL 10mM dNTPs, and 1 µL RT-MMVL enzyme (Promega) were added, and the mixture was incubated at 37°C for 30 minutes to generate cDNA.

### Fluidigm chip preparation

Gene expression analysis was conducted using a Fluidigm chip assay. Samples obtained post-reverse transcription were pre-amplified with Fluidigm products following the ‘Pre-amplification of cDNA for Gene Expression with delta GeneAssays’ protocol provided by the manufacturer (Biomark, Stan-dard Biotools Inc. San Francisco, CA, USA). Initially, 0.5 µL of 100 µM forward and reverse primers were diluted in 200 µL of DNA suspension buffer. A pre-mix was prepared by combining 105.6 µL of PreAmp Master Mix, 52.8 µL of the diluted primer mix, and 237.6 µL of DNase-free H_2_O. Each well of a 96-well plate was filled with 3.8 µL of the pre-mix and 1.75 µL of 1/10 diluted cDNA sample. The plate was incubated with the following program: 2 minutes at 95°C, followed by 10 cycles of 15 seconds at 95°C and 4 minutes at 60°C. Post pre-amplification, a clean-up reaction was performed by preparing an exonuclease mix (168 µL of DNase-free H_2_O, 24 µL of Exonuclease I Reaction Buffer, and 48 µL of Exonuclease I at 20 U/µL). Each well received 2 µL of this mix, and the plate was incubated for 30 minutes at 37°C for digestion, followed by 15 minutes at 80°C to inactivate the enzyme. Subsequently, 43 µL of DNA suspension buffer was added to each well, diluting the samples to one-tenth of their concentration, and the plate was stored at -20°C. For Fluidigm chip preparation, the “primers” and “samples” plates were arranged. The “primers” plate was prepared by mixing 3 µL of assay loading reagent, 2.7 µL of DNA suspension buffer, and 0.3 µL of 100 µM primers (forward + reverse). The “samples” plate mix contained 360 µL of SoFast EvaGreen Super Mix and 36 µL of DNA binding dye sample loading reagent, with 3.4 µL of this mix and 2.75 µL of pre-amplified samples added to each well, achieving a final volume of 5 µL per well. The software to obtain the data was Fluidigm Real-Time PCR Analysis, with the quality threshold set at 0.65. Gene expression for 95 genes, identified from the literature, was measured and normalized using the 2^ΔΔCt method (Livak & Schmittgen, 2001), with the 2 reference genes Tubulin and ATPase Ta54227, implemented using the R package FLUIDIGR. Genes with undetected expression (CT = 999) were excluded from the dataset. Any gene or sample with 20% or more missing data was removed from the analysis.

### Metabolomic analysis

LCMS-based untargeted metabolomics was performed from ethanol extracts as described previously (Berger *et al*. 2024). Briefly, 5mg of dried leaf and root metabolite extracts were extracted in 2 x 300 µL (ethanol/water 80% v/v, Formic acid 0.1% v/v, Methyl vanillate 250µg/mL), sonicated (15 min, 4°C), centrifuged (5 min, 10 600 g, 4°C) and filtered (sterile 0.22µm Durapore membrane, Merck), then analyzed by UHPLC-Q-Exactive Orbitrap MS/MS-Based Untargeted Metabolomics using a Vanquish ultra-high-pressure liquid chromatography (UHPLC) system coupled to a QEX+ mass spectrometer interfaced with an electrospray (ESI) ionisation source (ThermoScientific, Bremen, Germany). MS spectra were acquired in the negative ion mode at 35k resolution and LCMS/MS acquisitions (top N=2) were acquired at stepped normalized collision energies of 15, 30 and 40. The LCMS analytical sequence of 26 injections was randomized and included 18 unique biological samples (n = 3, 5 genotypes and 1 bulk soil control), 2 extraction blanks (prepared without biological material to rule out potential contaminants detected by untargeted metabolomics) and 6 Quality Control (QC) samples prepared by mixing 50 µL from each sample and biological standard. QC samples were used for i) the correction of signal drift during long batches, and ii) the calculation of coefficients of variation for each metabolomic feature so only the most robust ones are retained for chemometrics (Broadhurst et al., 2018). Raw LCMS data were processed following the DIA MS2 deconvolution method using MS-DIAL software (v. 4.9; (Tsugawa et al., 2015)) following optimized parameters. Annotations were performed based on MS1 spectra and MS2 DDA fragmentation information using FragHUB database, including thousands of phytochemicals (Dablanc *et al*. 2024). Thus, putative annotation of differentially accumulated metabolites resulted from MS-DIAL screening of the MS1 detected exact HR m/z and MS2 fragmentation patterns (Tsugawa et al., 2015). Additionally, the InChiKeys of annotated features were employed within ClassyFire to generate a structural ontology for chemical entities (Djoumbou Feunang et al., 2016).

Curation of 6369 raw metabolomic signals (SN > 10, CV QC < 30%) resulted in 1168 LCMS features, the intensities of which were standardized by the mass of the dry extracts. Among these, 1104 remained unidentified, having no match with either MS1 or MS2. Another 32 features were suggested to be annotated metabolites, having a match with MS1 only (referred to as level 3 MS ID), while 64 features were positively annotated as metabolites, matching both MS1 and MS2 (referred to as level 2 MS ID). These 1168 features were normalised (Pareto scaling) for normality and comparison purposes, and underwent uni- and multivariate statistical analyses (Principal Component and Clustering analyses) using MetaboAnalyst (v 6) (Pang et al., 2021).

### Data analyses

All statistical analyses were performed in R v.4.3.1 (R Core Team, 2023).

#### Phenotypic analyses

Boxplots were created to visualize STB symptoms, aerial and root phenotypes using the ggplot2, see, and stats packages. To evaluate the blend effect (pure *vs*. mixed), contrast tests were conducted following an aligned and ranked linear model with the ARTool package (Equation 1):

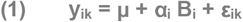

where y_ik_ is the phenotypic observation; μ is the intercept; α_i_ corresponds to the effect for the blend type i; B_i_ corresponds to the blend type i; and ε_ik_ is the error term.

Root phenotyping differences between Cultur and Atoudur under mixed conditions were further assessed with non-parametric contrast tests.

#### Transcriptional analyses

Transcriptional profiles were visualized using Principal Component Analysis (PCA) with the factomineR and factoextra packages, focusing on leaf transcriptional profiles at 10 and 21 days after germination (dag) and defense gene expression at 7 and 14 days post inoculation (dpi). Boxplots of log2 fold-changes in the expression of 14 defense-related genes in Cultur leaves grown in pure stands *versus* mixed with Atoudur at 7 and 14 dpi, and barplots of fold-changes in gene expression under mixed conditions compared to pure stands at 10 and 21 dag were generated using the ggplot2, see, and stats packages. The effects of blend conditions on PCA coordinates and gene expression at 10 and 21 dag were analyzed with non-parametric ANOVA followed by Benjamini-Hochberg correction (Equation 1). Additionally, log2 fold-changes for defense-related genes were compared between pure and mixed conditions using Wilcoxon tests.

#### Metabolomic analyses

The analyses were performed on the metabolites with the highest confidence level in metabolite identification based on the LC-MS profile of authentic pure standards, including retention time, m/z ratio, and fragmentation spectra when available (level 1 identification).

Metabolomic profiles were visualized using heatmaps (gplots package) and Principal Component Analysis (PCA) (factomineR and factoextra packages). Heatmaps were generated for leaf and root metabolomic data at 21 dag and 7 dpi. Hierarchical clustering for heatmaps was conducted using the complete clustering method with Euclidean distance. PCA was performed on leaf and root metabolites at 21 dag. The effects of blend conditions on PCA coordinates were analyzed with non-parametric ANOVA (Equation 1).

For metabolite analysis at 21 dag, log2 fold-changes between the mixed and pure conditions and z-scores (Equation 2) were calculated. Metabolites were considered significantly differentially accumulated if they showed a log2 fold-change of less than -1 or greater than 1 and a z-score below -2 or above 2.

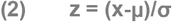

where z is the vector of z-scores; x is the log2 fold-change of metabolite accumulation between mixture and pure condition; μ is the mean log2 fold-changes; and σ is the standard error of the log2 fold-changes.

For metabolite analysis at 7 dpi, a similar approach was used, with log2 fold-changes calculated between inoculated and mock-treated conditions. This comparison was conducted separately in pure and mixed conditions.

## Results

### The neighbor genotype Atoudur increases Septoria tritici blotch susceptibility and impacts phenological traits in the genotype Cultur

As previously reported (Pélissier *et al*. 2021a), Cultur was more susceptible to STB when grown with Atoudur than in pure stands (Fig. 1A). To assess whether traits other than STB susceptibility are influenced by the Cultur-Atoudur interaction, we examined aerial and root traits of Cultur in pure and mixed conditions. Chlorophyll content was decreased in Cultur leaves in mixture, in contrast to plant height, and leaf or tiller number (Fig. 1B and Supplementary Fig. S3A). Root traits remained unchanged in Cultur grown in pure stand or mixed with Atoudur at the seedling stage (Supplementary Fig. S3B). However, at 8 weeks after germination, the flowering rate was higher in Cultur grown with Atoudur than in pure (Fig. 1C), indicating a mixture effect on subsequent developmental stages. The results show that plant-plant interactions specifically impact STB susceptibility, leaf chlorophyll content and heading date.

**Fig. 1.**
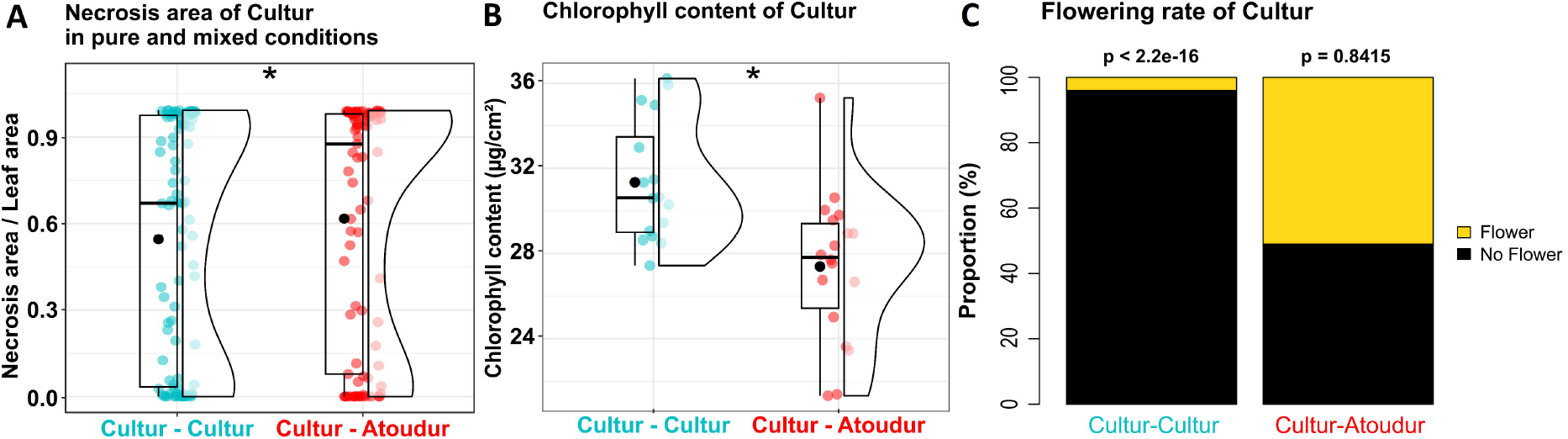
Effects of neighbor genotype on Septoria tritici blotch symptoms, chlorophyll content and flowering rate in a durum wheat cultivar mixture. **A** Septoria tritici blotch symptoms measured as the part of necrotic leaf area 17 days after inoculation and **B** chlorophyll content assessed 38 days after germination, both on the genotype Cultur grown either in pure stands or in mixtures with Atoudur. Statistical comparisons between Cultur in pure and mixed conditions were conducted using non-parametric contrast tests. Significance levels are denoted as follows:.: p < 0.1, *: p < 0.05, **: p < 0.01, ***: p < 0.001. **C** The flowering rate of the genotype Cultur assessed 56 days after germination. χ^2^ test showed significantly higher flowering in Cultur when grown with the neighbor genotype Atoudur.

### The increased STB susceptibility of the genotype Cultur in mixture is driven by neighbor root proximity independently of phenolic exudates

We tested several hypotheses related to root-based mechanisms potentially involved in plant-plant interactions: enhanced susceptibility in the mixture could be due either to differential soil metabolite contents or to differential competition in the soil. First, we introduced a porous barrier between genotypes to hamper direct root contact. Under these conditions, no significant difference in STB symptoms was observed between pure and mixed conditions (Fig. 2A) in contrast with no barrier (Fig. 1A). Similarly, when the barrier was filled with polyvinylpolypyrrolidone (PVPP) to block the exchange of phenolics (Durán-Lara *et al*. 2015), STB susceptibility remained unchanged between pure and mixed conditions (Fig. 2A). Therefore, physical impediment of root-root contacts is sufficient to suppress the neighbor-mediated increase in susceptibility.

**Fig. 2.**
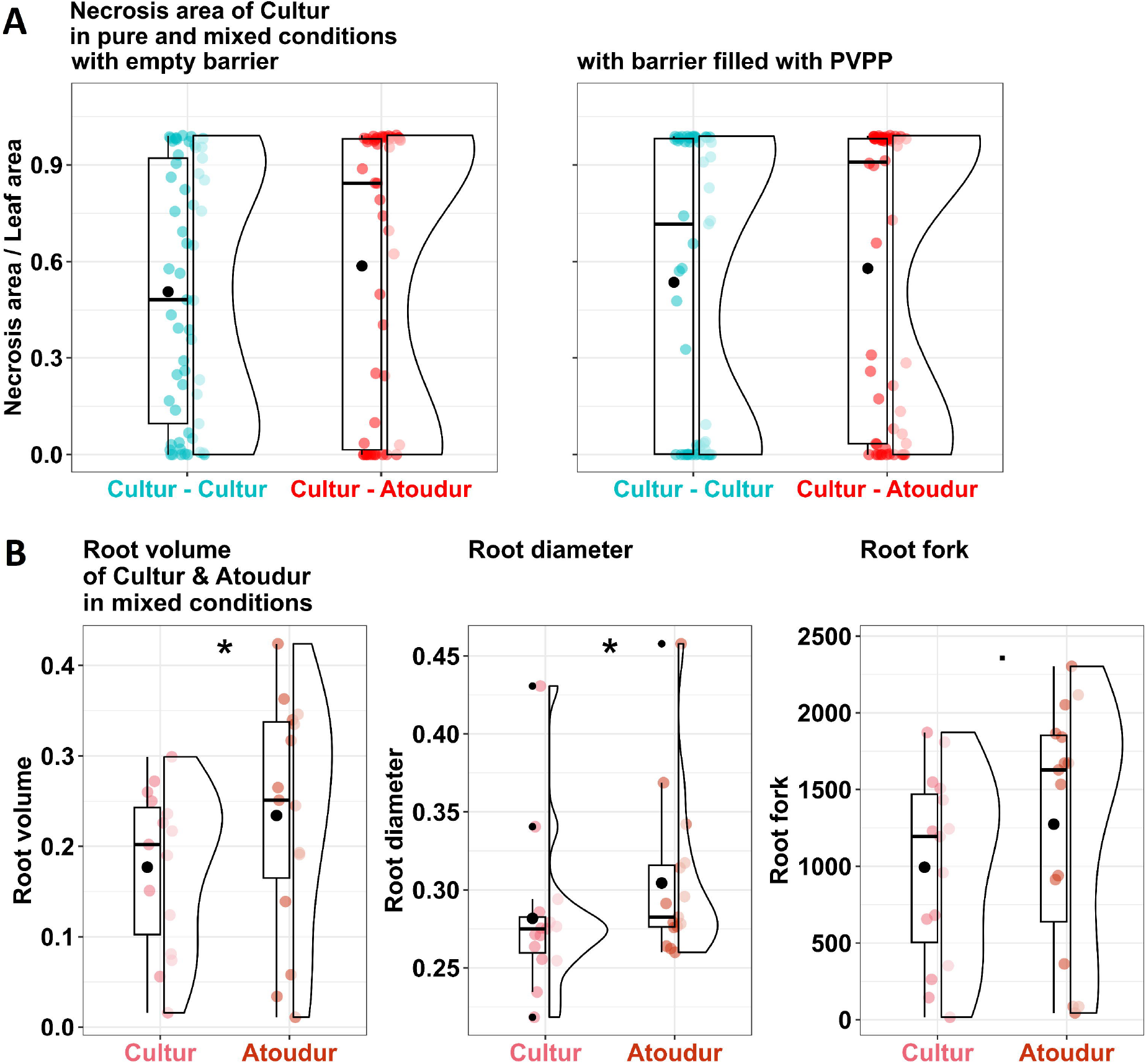
Effects of neighbor genotype and root barrier on Septoria tritici blotch symptoms, and assessment of root traits in a durum wheat cultivar mixture. **A** Septoria tritici blotch symptoms of the genotype Cultur were assessed by necrotic leaf area 17 days after inoculation. Cultur was grown in pure stands or mixed with Atoudur, with either an empty barrier or a barrier filled with polyvinylpolypyrrolidone (PVPP) between genotypes. No statistical difference between Cultur symptoms in pure and mixed conditions were observed based on non-parametric contrast tests. **B** Root phenotyping of Cultur and Atoudur in mixed conditions, measured 21 days after germination (day of inoculation). Statistical comparisons between Cultur and Atoudur phenotypes were conducted using non-parametric contrast tests. Significance levels are denoted as follows:.: p < 0.1, *: p < 0.05, **: p < 0.01, ***: p < 0.001.

Second, we measured various root traits in mixtures to evaluate if obvious root architecture could be responsible for differential competition. Interestingly, this analysis revealed differences between the two genotypes, with Atoudur displaying a significantly more prominent root system than Cultur, including root volume, root diameter and root branching (Fig. 2B). Altogether, these results are in line with the idea that Cultur trait changes are induced by close root-root contacts with Atoudur.

### A metabolic slow-down in Cultur when co-cultivated with Atoudur

We next addressed the question of the molecular responses in the focal plant using metabolomic and transcriptional approaches, in both roots and leaves, in the absence of infection.

In the roots, a significant effect was detected in the metabolomic profiles of Cultur (Fig. 3A). We observed a small cluster of slightly up-accumulated metabolites in the mixture, including two phenylpropanoids (cluster R1 in Supplementary Fig. S4A, Supplementary Fig. S5, Supplementary Table S2). In contrast, there was a large cluster of metabolites that were tendentially down-regulated (cluster R2 in Supplementary Fig. S4A, Supplementary Fig. S6, Supplementary Table S2). This cluster was enriched in metabolites involved in primary metabolism pathways (75% of metabolites present in this cluster; Supplementary Table S2), even though the strongest down-regulations were observed for phenylpropanoids. Thus, the presence of Atoudur roots in the neighborhood of Cultur has a general negative impact on metabolic activity in Cultur roots.

**Fig. 3.**
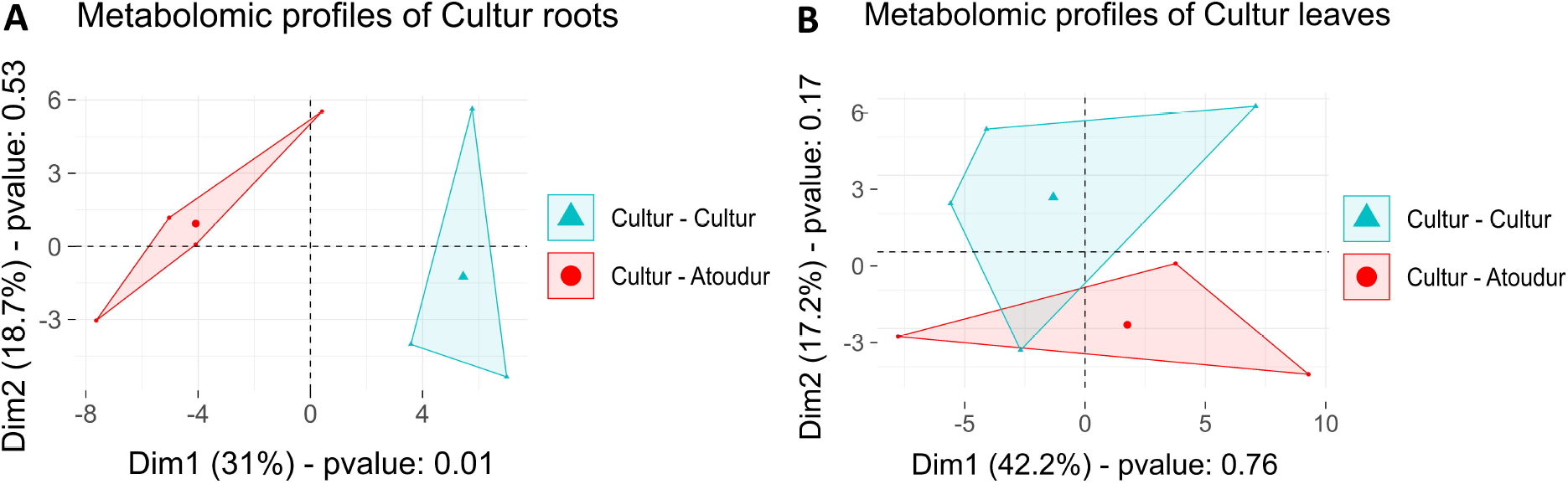
Effects of neighbor genotype on metabolomic profiles in a durum wheat cultivar mixture. **A**,**B** Principal component analysis (PCA) of **A** root and **B** leaf metabolomic profiles from Cultur grown in pure stands or mixed with Atoudur, sampled 21 days after germination (day of inoculation). Statistical analyses between pure and mixed conditions for each PCA dimension were performed using a non-parametric ANOVA.

In the leaves, a slight effect of the mixture was observed in the leaf metabolomic profiles of Cultur (Fig. 3B). In Cultur leaves, we identified a cluster of mildly up-accumulated metabolites related to specialized metabolism (43% annotated as phenylpropanoids) in the mixture (cluster L2 in Supplementary Fig. S4B, Supplementary Fig. S7, Supplementary Table S2). Conversely, a separate cluster of down-accumulated metabolites was observed (cluster L1 in Supplementary Fig. S4B, Supplementary Fig. S8, Supplementary Table S2), with 8 metabolites related to primary metabolism among the 12 metabolites also down-regulated in mixed roots (cluster R2 in Supplementary Fig. S4A, Supplementary Fig. S6, Supplementary Table S2).

While these metabolic changes in the leaves were slight, significant global changes were seen in the transcriptional profiles of Cultur both at 10 days after germination (Fig. 4A) and at 21 dag (before inoculation) (Fig. 4B). In particular, at 10 dag, there was a down-regulation of *SS4* and *SHMT* genes used as markers for carbon metabolism (Fig. 4C and Supplementary Fig. S9), and at 21 dag, a down-regulation of marker genes for cell expansion (*PIF3*), potassium (*HAK25*) and nitrogen (*AMT1*.*1*) metabolisms (Fig. 4D and Supplementary Fig. S10). These changes were consistent with metabolic data from Cultur leaves at 21dag showing a reduction of metabolites associated with primary metabolism (Supplementary Fig. S4B).

**Fig. 4.**
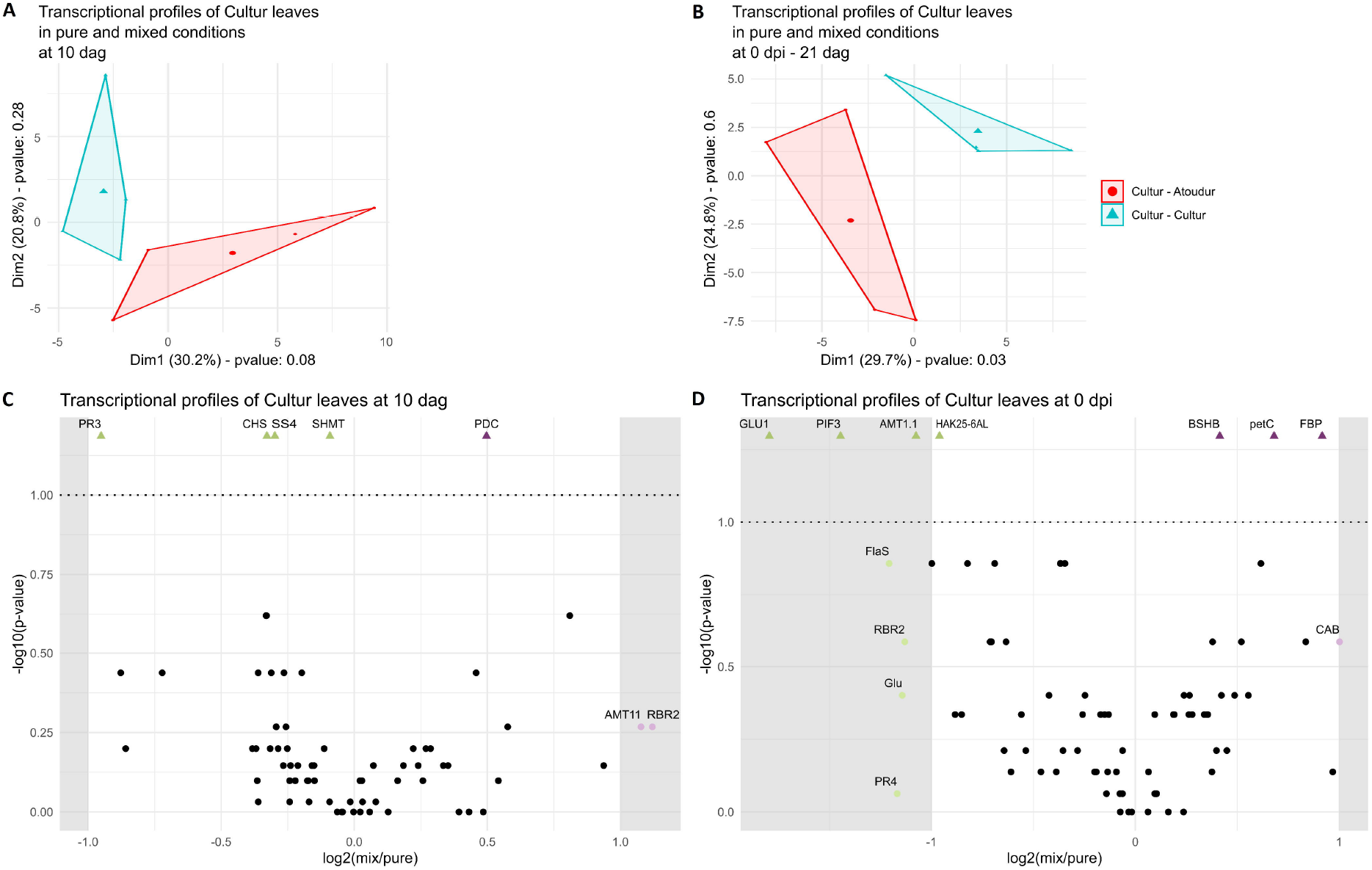
Effects of neighbor genotype on transcriptional profiles in a durum wheat cultivar mixture. **A**,**B** Principal component analysis (PCA) of transcriptional profiles from Cultur leaves grown in pure stands or mixed with Atoudur, sampled **A** 10 days after germination or **B** 21 days after germination (day of inoculation). Statistical analyses between pure and mixed conditions for each PCA dimension were performed using a non-parametric ANOVA. **C**,**D** Scatterplots showing log2 fold-changes in gene expression of Cultur between mixed and pure conditions *versus* the -log10 p-value for each gene, with samples collected at **C** 10 days after germination and **D** 21 days after germination (day of inoculation). Statistical analysis was performed using a non-parametric ANOVA followed by Benjamini-Hochberg correction to compare Cultur gene expression in pure *vs*. mixed conditions for each gene.

Altogether, these analyses in roots and leaves indicate that, as early as 10 days of co-culture, interactions at the root level resulted in detectable changes at the molecular level in the leaves. However, with the exception of the *PR3* gene, no massive change in the expression of 15 tested defense genes could be detected before infection. In fact, the changes observed were more global and reflective of an induced resource scarcity at the whole organismic level in the Cultur genotype when grown nearby the competitive Atoudur genotype.

### Delayed induction of defense gene expression and reduced accumulation of specialized metabolism compounds in Cultur grown with Atoudur

To analyze the impact of the Cultur-Atoudur interaction on the response to *Zymoseptoria tritici* inoculation, we compared leaf expression of induced defense genes between inoculated and mock-treated Cultur in pure and mixed conditions. This analysis was conducted at two time points: 7 and 14 days post-inoculation (dpi), corresponding to the biotrophic and necrotrophic phases of pathogen development, respectively.

At 7 dpi, Cultur in pure stands showed distinct defense responses between mock and inoculated leaves, whereas no difference was observed in mixed conditions (Fig. 5A). This divergence between mixed and pure conditions was linked to an absence of induction of defense genes in mixed conditions compared to a two-fold induction in pure conditions (Fig. 5B). By 14 dpi, however, the defense response was similar between pure and mixed conditions, with no further difference in gene expression (Fig. 5C & 5D). These findings suggest a delay in defense gene induction in response to STB inoculation when Cultur is grown with Atoudur.

**Fig. 5.**
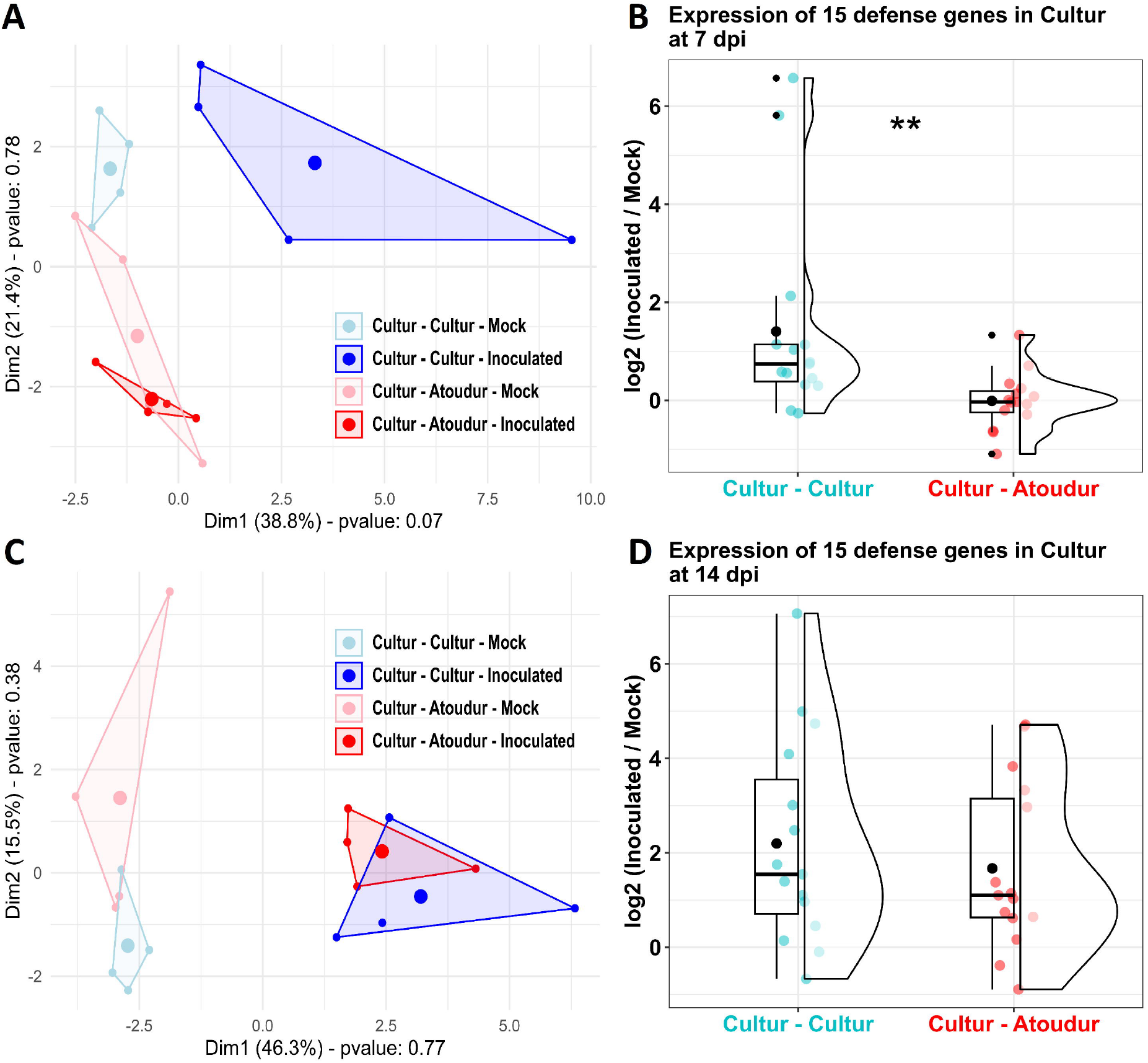
Defense gene expression in mock-treated and inoculated focal leaves grown in pure stands *vs*. in a durum wheat cultivar mixture. **A**,**C** Principal component analysis (PCA) of defense gene expression profiles in Cultur leaves at **A** 7 days and **C** 14 days after inoculation, comparing inoculated and mock-treated leaves grown in either pure Cultur stands or mixed with Atoudur. **B**,**D** Log2 fold-changes in expression of 15 defense genes between inoculated and mock-treated Cultur leaves grown in pure stands *versus* mixed with Atoudur at **B** 7 days and **D** 14 days after inoculation. The defense genes are listed in Supplementary Table S1. Statistical analyses between pure and mixed conditions were performed using a Wilcoxon test. Significance levels are denoted as follows:.: p < 0.1, *: p < 0.05, **: p < 0.01, ***: p < 0.001.

We also examined metabolites at 7 dpi and observed that 60% of regulated metabolites in response to inoculation under pure condition showed no difference in the mixed condition. A notable cluster of specialized metabolism compounds, including triterpenoids, and precursors of phenolics and alkaloids, displayed this pattern similar to defense genes at 7 dpi (Fig. 6, Supplementary Table S2).

**Fig. 6.**
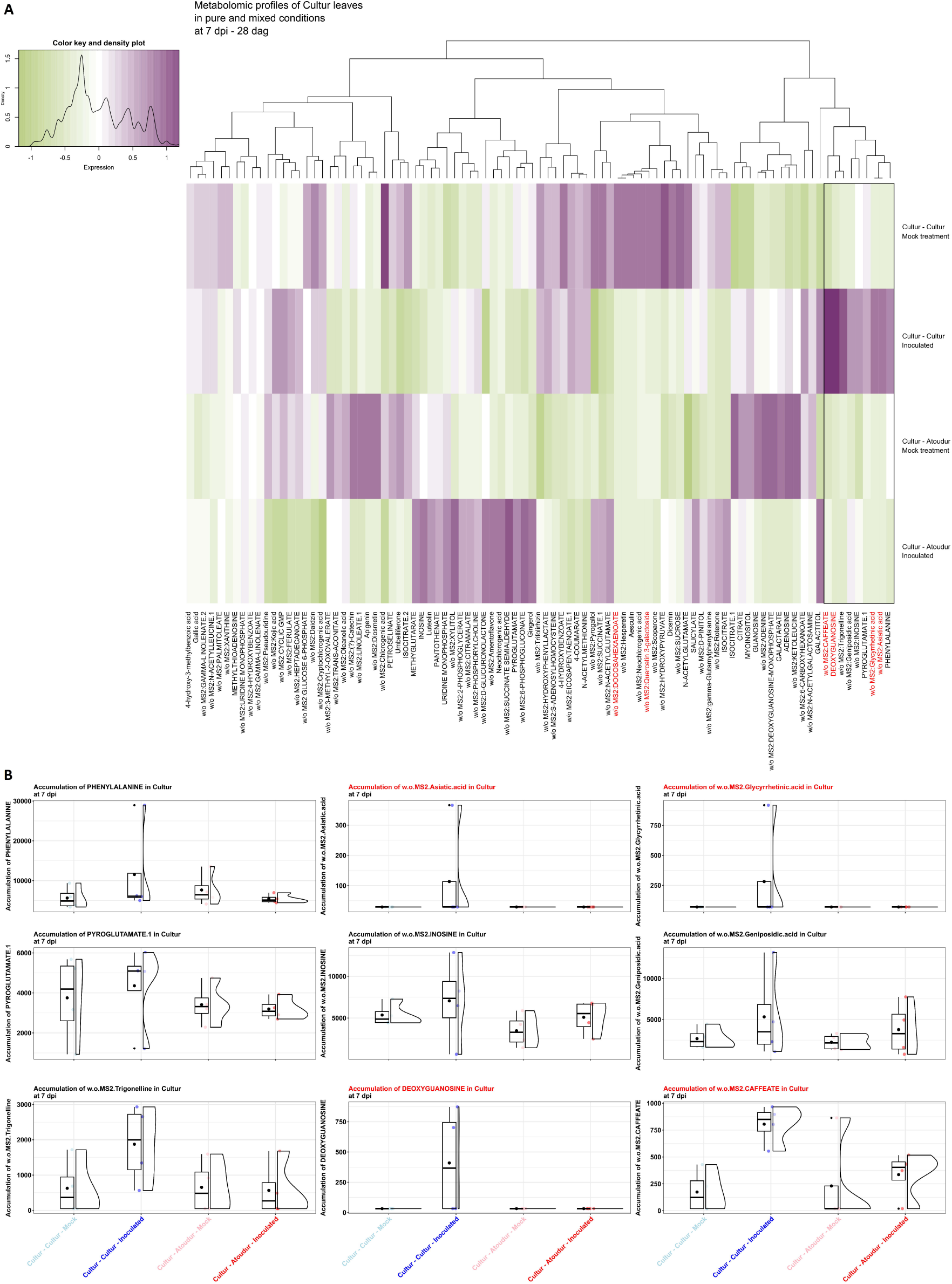
Metabolomic profiles of focal leaves grown in pure stands *vs*. in a durum wheat cultivar mixture under mock-treated and inoculated conditions. **A** Heatmap of metabolomic profiles in Cultur leaves, grown in pure stands *versus* in a mixture with Atoudur, analyzed at 7 days post inoculation. **B** Boxplots of metabolites showing differences in Cultur metabolite accumulation between inoculated and mock-treated conditions in pure stands and not in the mixed condition. Metabolites meeting the significance criteria (log2 fold-change < -1 or > 1 and z-score < -2 or > 2 in pure stands, with log2 fold-change > -1 or < 1 and z-score > -2 or < 2 in mixture) are marked in red to indicate significant differential accumulation only in pure stands.

## Discussion

This study on the enhanced susceptibility of the durum wheat variety Cultur to Septoria tritici blotch (STB) in mixed stands with Atoudur reinforces the role of root-mediated interactions in increasing STB severity of Cultur. Our findings align with previous work by (Pélissier *et al*. 2021a), who observed similar belowground effects influencing susceptibility to leaf rust. To further explore whether this root-based signal involves resources or signaling molecules, we used a porous barrier that permitted nutrient and small molecule diffusion while decreasing physical root contacts. Our results suggest that the increased susceptibility in Cultur requires close physical root contacts and/or unequal spatial occupation and/or diffusible molecules at short distance. Differences in root architecture between Cultur and Atoudur, with the latter showing a more developed root system, suggest that resource competition could be the trigger of the observed phenotypes. Indeed, particularly under nutrient-limited conditions, competition for nutrients, water and space plays a critical role in plant fitness, including productivity and development (Schenk 2006).

The analysis of the molecular responses in the focal plants Cultur provided hints into the processes that the neighboring genotype Atoudur induced. Indeed, our transcriptional and metabolomic analyses revealed shifts in metabolic pathways towards a metabolic shutdown in roots, characterized by a downregulation of primary metabolism. The negative regulation of carbon metabolism and cell expansion pathways, along with the observed reduction of chlorophyll content, could suggest a competition-induced inhibition of photosynthesis-related pathways, consistent with previous studies (Schmidt & Baldwin 2006; Horvath, Gulden & Clay 2006; Masclaux *et al*. 2012; Pierik, Mommer & Voesenek 2013). On the other hand, the increase in phenylpropanoids, known to be involved in defense and stress responses (Chowdhary, Alooparampil, V. Pandya & G. Tank 2022), further indicates that Atoudur activates stress-related pathways in Cultur. Similar patterns, including the up-regulation of flavonoids (Bowsher *et al*. 2017), were observed in response to competitive interactions (Schmidt & Baldwin 2006; Masclaux *et al*. 2012). Overall, the molecular shifts in the Cultur-Atoudur mixture likely reflect a response to competition, preceding any later observable phenotypic changes. For instance, early flowering, a well-documented response that enables increased plant survival in stressful environments (Takeno 2016), was observed in the Cultur-Atoudur mixture. This change in phenology is also a hallmark of competition in plants (Moreno-Colom & Montesinos-Navarro 2025) In that respect, the mis-regulation of the *PIF3* gene in Cultur grown with Atoudur is consistent with the regulation of reproduction, as observed in Arabidopsis (Galvāo *et al*. 2019).

The increase in STB susceptibility in Cultur mixed with Atoudur was associated with a delay in the activation of defense genes and the accumulation of specialized metabolites, including terpenoids and precursors of phenolics and alkaloids, particularly at the biotrophic stage. The delay in defense response is important, as early defense mechanisms play a key role in inhibiting disease progression (Brennan *et al*. 2019). Furthermore, (Seybold *et al*. 2020) reported a delayed accumulation of phenylpropanoid compounds and flavonoids in the susceptible wheat cultivar compared to the resistant one when exposed to *Z. tritici*. This suggests that these compounds, known for their antimicrobial and antioxidant properties (Treutter 2006), are essential in resistance to STB infection. Therefore, the compromised defense gene expression and immune-related metabolites observed in Cultur grown with Atoudur likely facilitated pathogen activity, resulting in increased STB severity. It is tempting to speculate that this compromised defense resulted from a trade-off between growth and defense triggered by the abovementioned competition. However, this remains to be demonstrated.

Our integrated approach, combining aerial and root phenotyping, transcriptional analysis, and the first untargeted metabolomics in a varietal mixture, provides new insights into root-mediated mechanisms influencing disease susceptibility. The findings indicate that the increased STB severity observed in Cultur when co-cultivated with Atoudur could be primarily driven by root architecture differences and resource competition rather than allelochemical signaling (Fig. 7). While Atoudur’s large roots may induce early molecular changes before pathogen inoculation in Cultur, these changes likely reflect competitive stress and potentially impact the phenology of Cultur. Competition and the subsequent reduction of primary metabolism may indirectly reduce the expression of processes like defenses, thus explaining the observed delay in defense responses (Fig. 7). These results highlight the interplay between root architecture, resource competition, plant metabolism, and defense response in modulating plant-pathogen interactions in varietal mixtures. Future research should focus on disentangling these root-mediated mechanisms and assessing their potential in enhancing crop resilience in mixtures.

**Fig. 7.**
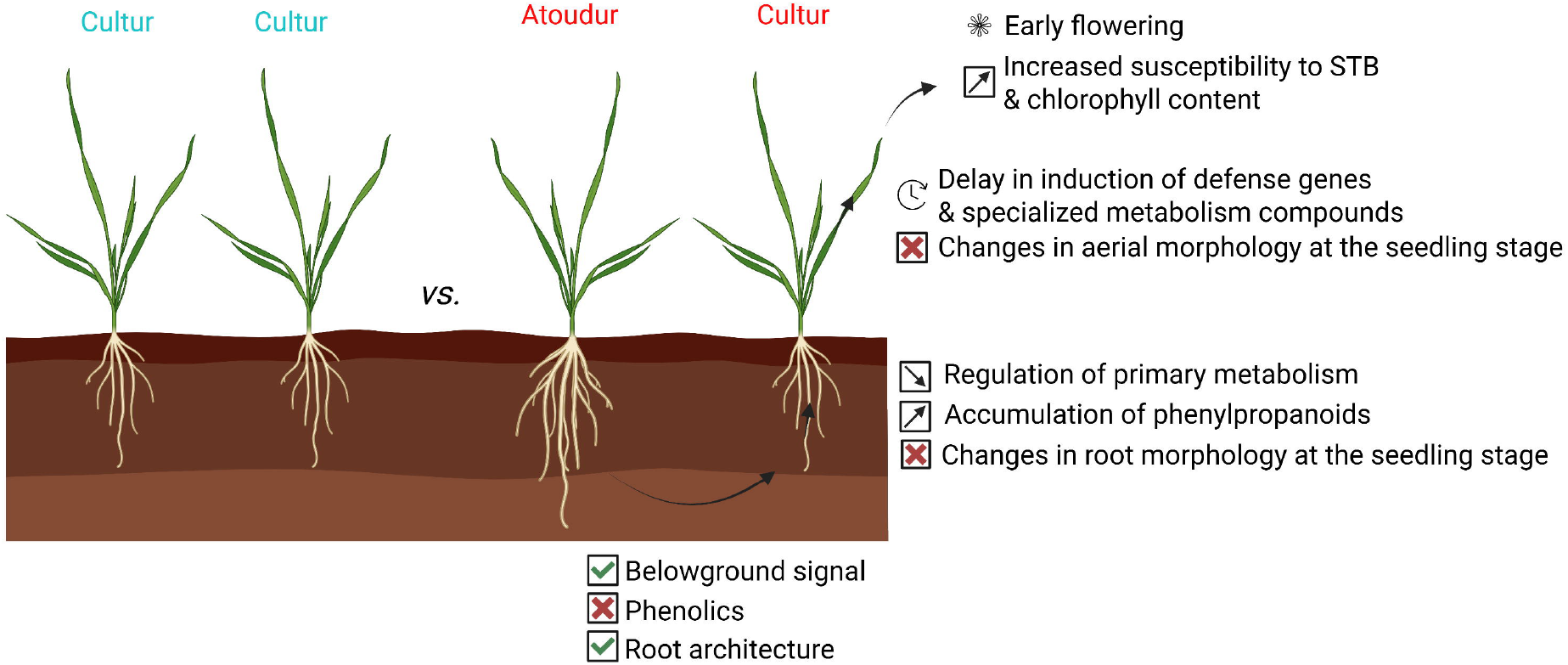
Scheme of the proposed signaling cascade in a durum wheat varietal mixture. In the Cultur-Atoudur mixture, a belowground signal—presumably associated with root architecture rather than root-derived metabolites—triggers a competitive response characterized by the down-regulation of primary metabolism and an increase in phenylpropanoid accumulation in Cultur. Additionally, the delayed activation of defense following Septoria tritici blotch (STB) inoculation likely explain the observed higher susceptibility to STB in this varietal mixture.

## Supplementary data

The following supplementary data are available at JXB online.

Supplementary Fig. S1. Experimental design for studying the modulation of *Zymoseptoria tritici* susceptibility in a durum wheat cultivar mixture.

Supplementary Fig. S2. Overview of biological processes represented in the high-throughput RT-qPCR chip for wheat transcriptional response analysis.

Supplementary Fig. S3. Effect of neighbor genotype on aerial and root traits in a durum wheat cultivar mixture.

Supplementary Fig. S4. Effects of neighbor genotype on metabolomic profiles in a durum wheat cultivar mixture.

Supplementary Fig. S5. Accumulation profiles of metabolites of cluster “R1” in Cultur roots grown in pure stands *vs*. in a durum wheat cultivar mixture.

Supplementary Fig. S6. Accumulation profiles of metabolites of cluster “R2” in Cultur roots grown in pure stands *vs*. in a durum wheat cultivar mixture.

Supplementary Fig. S7. Accumulation profiles of metabolites of cluster “L2” in Cultur leaves grown in pure stands *vs*. in a durum wheat cultivar mixture.

Supplementary Fig. S8. Accumulation profiles of metabolites of cluster “L1” in Cultur leaves grown in pure stands *vs*. in a durum wheat cultivar mixture.

Supplementary Fig. S9. Comparative analysis of gene expression grouped by biological processes in focal leaves grown in pure stands *vs*. in a durum wheat cultivar mixture at 10 days after germination.

Supplementary Fig. S10. Comparative analysis of gene expression grouped by biological processes in focal leaves grown in pure stands *vs*. in a durum wheat cultivar mixture at 21 days after germination.

Supplementary Table S1. Detailed list of genes included in the high-throughput RT-qPCR chip for analyzing transcriptional responses in wheat (*Triticum aestivum* and *Triticum turgidum*).

Supplementary Table S2. Detailed list of detected metabolites.

## Acknowledgments

We kindly thank Sandrine Roques for providing the rhizoboxes, and Christophe Jourdan and Didier Arnal for the loan of the WinRhizo system used for root phenotyping. We also thank Aurélie Ducasse, Luis Buendia and Andy Brousse for producing preliminary data that are not shown here but helped us in designing later experiments.

## Author contributions

LM conceived and designed the experiments. LM, AC, and SM performed aerial phenotyping. LM, AC, and JS conducted root phenotyping. LM and AC performed the experiments and RNA extractions. CR, JV, and PP generated the metabolomic data. LM conducted all computational analyses and prepared the figures and tables. LM led the writing of the manuscript. JBM, LVM, and EB supervised the work. LM, JBM, LVM, and EB reviewed the manuscript. All authors read and approved the final version.

## Data availability

The data that support the findings of this study are openly available in Zenodo at doi.org/10.5281/zenodo.15600332.

## Figure legends

**Supplementary Fig. S1.** Experimental design for studying the modulation of *Zymoseptoria tritici* susceptibility in a durum wheat cultivar mixture. 10 days after germination: Transcriptional analysis was performed on leaves of Cultur to compare gene expression profiles between plants grown in pure stands and those mixed with Atoudur. 21 days after germination (day of inoculation): Transcriptional and metabolomic analyses were conducted on both leaves and roots of Cultur to assess differences between pure stands and mixtures with Atoudur. Metabolomic analysis of soil associated with Cultur was performed, comparing samples from pure stands and mixtures with Atoudur under control conditions or with a barrier containing polyvinylpolypyrrolidone (PVPP) between genotypes. Root phenotyping was carried out to compare root architecture between Cultur and Atoudur under mixed conditions. 7 days post inoculation (biotrophic phase): Transcriptional and metabolomic analyses were conducted on Cultur leaves to compare responses between inoculated and mock-treated leaves grown in pure stands *versus* mixed with Atoudur. 14 days post inoculation (necrotrophic phase): Transcriptional analysis of Cultur leaves was performed to evaluate differences in gene expression between inoculated and mock-treated plants grown in pure *versus* mixed conditions. 17 days post inoculation: Aerial phenotyping and assessment of disease symptoms were conducted on Cultur to compare the level of *Zymoseptoria tritici* susceptibility between plants grown in pure stands *versus* mixed with Atoudur.

**Supplementary Fig. S2.** Overview of biological processes represented in the high-throughput RT-qPCR chip for wheat transcriptional response analysis. The RT-qPCR chip is designed to assess the transcriptional responses in wheat by targeting a comprehensive selection of marker genes across multiple biological processes. The chip includes genes associated with defense (15 genes), carbon metabolism (10), nitrogen metabolism (9), cell state (8), housekeeping functions (5), flavonoid biosynthesis (5), cytokinin signaling (4), photosynthesis (2), phosphate metabolism (3), potassium regulation (3), jasmonic acid signaling (3), gibberellic acid signaling (3), ethylene signaling (3), drought response (3), DIMBOA biosynthesis (3), brassinosteroid signaling (3), auxin signaling (3), abscisic acid signaling (3), strigolactones (2), shade avoidance syndrome (3), and salicylic acid signaling (2). The full list of marker genes is detailed in Supplementary Table S1.

**Supplementary Fig. S3.** Effect of neighbor genotype on aerial and root traits in a durum wheat cultivar mixture. **A** Aerial and **B** root phenotyping of the genotype Cultur was assessed 38 and 21 days after germination, respectively. Plants were grown either in pure stands or in mixtures with Atoudur. No statistical differences in aerial and root traits were observed between Cultur grown in pure stands and mixed conditions, based on non-parametric contrast tests.

**Supplementary Fig. S4.** Effects of neighbor genotype on metabolomic profiles in a durum wheat cultivar mixture. **A**,**B** Heatmap of Cultur metabolite accumulation in **A** roots and **B** leaves, grown either in pure stands or mixed with Atoudur, also analyzed at 21 days after germination (day of inoculation). Metabolites meeting the significance threshold (log2 fold-change < -1 or > 1 and z-score < -2 or > 2) are marked in red to indicate significant differential accumulation.

**Supplementary Fig. S5.** Accumulation profiles of metabolites of cluster “R1” in Cultur roots grown in pure stands *vs*. in a durum wheat cultivar mixture. Boxplots of Cultur metabolite accumulation between pure stands and mixed with Atoudur at 21 days after germination (day of inoculation). Metabolites meeting the significance criteria (log2 fold-change < -1 or > 1 and z-score < -2 or > 2) are marked in red to indicate significant differential accumulation.

**Supplementary Fig. S6.** Accumulation profiles of metabolites of cluster “R2” in Cultur roots grown in pure stands *vs*. in a durum wheat cultivar mixture. Boxplots of Cultur metabolite accumulation between pure stands and mixed with Atoudur at 21 days after germination (day of inoculation). Metabolites meeting the significance criteria (log2 fold-change < -1 or > 1 and z-score < -2 or > 2) are marked in red to indicate significant differential accumulation.

**Supplementary Fig. S7.** Accumulation profiles of metabolites of cluster “L2” in Cultur leaves grown in pure stands *vs*. in a durum wheat cultivar mixture. Boxplots of Cultur metabolite accumulation between pure stands and mixed with Atoudur at 21 days after germination (day of inoculation). Metabolites meeting the significance criteria (log2 fold-change < -1 or > 1 and z-score < -2 or > 2) are marked in red to indicate significant differential accumulation.

**Supplementary Fig. S8.** Accumulation profiles of metabolites of cluster “L1” in Cultur leaves grown in pure stands *vs*. in a durum wheat cultivar mixture. Boxplots of Cultur metabolite accumulation between pure stands and mixed with Atoudur at 21 days after germination (day of inoculation). Metabolites meeting the significance criteria (log2 fold-change < -1 or > 1 and z-score < -2 or > 2) are marked in red to indicate significant differential accumulation.

**Supplementary Fig. S9.** Comparative analysis of gene expression grouped by biological processes in focal leaves grown in pure stands *vs*. in a durum wheat cultivar mixture at 10 days after germination. **A** Bar plots of the fold-change in gene expression of Cultur leaves grown in a mixture with Atoudur compared to pure stands at 10 days after germination. Genes are categorized by their associated biological processes. Statistical analysis was performed using non-parametric ANOVA followed by Benjamini-Hochberg correction to compare Cultur gene expression in pure *vs*. mixed conditions for each gene. Genes exhibiting twofold and/or significant differences are colored in dark, and genes with significant expression differences (adjusted p-value < 0.05) are labeled on the bar plots. The abbreviations of genes and biological processes, and their full names, are provided in Supplementary Supplementary Table S1. **B** Boxplots of genes showing significant differences of Cultur gene expression in pure *vs*. mixed conditions at 10 days after germination. Statistical analysis was performed using non-parametric ANOVA followed by Benjamini-Hochberg correction to compare Cultur gene expression in pure *vs*. mixed conditions for each gene. Significance levels are denoted as follows:.: p < 0.1, *: p < 0.05, **: p < 0.01, ***: p < 0.001.

**Supplementary Fig. S10.** Comparative analysis of gene expression grouped by biological processes in focal leaves grown in pure stands *vs*. in a durum wheat cultivar mixture at 21 days after germination. **A** Bar plots of the fold change in gene expression of Cultur leaves grown in a mixture with Atoudur compared to pure stands at 21 days after germination (day of inoculation). Genes are categorized by their associated biological processes. Statistical analysis was performed using non-parametric ANOVA followed by Benjamini-Hochberg correction to compare Cultur gene expression in pure *vs*. mixed conditions for each gene. Genes exhibiting twofold and/or significant differences are colored in dark, and genes with significant expression differences (adjusted p-value < 0.05) are labeled on the bar plots. The abbreviations of genes and biological processes, and their full names, are provided in Supplementary Supplementary Table S1. **B** Boxplots of genes showing significant differences of Cultur gene expression in pure *vs*. mixed conditions at 21 days after germination. Statistical analysis was performed using non-parametric ANOVA followed by Benjamini-Hochberg correction to compare Cultur gene expression in pure *vs*. mixed conditions for each gene. Significance levels are denoted as follows:.: p < 0.1, *: p < 0.05, **: p < 0.01, ***: p < 0.001.

**Supplementary Table S1.** Detailed list of genes included in the high-throughput RT-qPCR chip for analyzing transcriptional responses in wheat (*Triticum aestivum* and *Triticum turgidum*). This chip is designed to capture a broad spectrum of biological processes by incorporating marker genes involved in defense mechanisms, carbon and nitrogen metabolism, cell state regulation, housekeeping functions, phytohormone signaling (cytokinin, jasmonic acid, gibberellic acid, ethylene, abscisic acid, brassinosteroids, auxin, and strigolactones), as well as processes such as flavonoid biosynthesis, phosphate and potassium metabolism, photosynthesis, drought response, DIMBOA biosynthesis, and shade avoidance syndrome. For each gene, the table provides the gene name, full description, associated biological process, specific sub-process, gene IDs in both *T. aestivum* and *T. turgidum*, reference publication, and the expected transcriptional response within the analyzed processes.

**Supplementary Table S2.** Detailed list of detected metabolites. For each metabolite, the table provides the name, superclass, class and subclass, along with the pathways it is involved in, its KEGG ID, and RefMetaDB ID. Metabolites identified within specific clusters are indicated.

